# Disease mechanism, drug-target and biomarker prediction software: Application on prostate cancer and validation

**DOI:** 10.1101/129742

**Authors:** Gokmen Altay, Elmar Nurmemmedov, Santosh Kesari, David E. Neal

## Abstract

We present an R software package that performs at genome-wide level differential network analysis and infers only disease-specific molecular interactions between two different cell conditions. This helps revealing the disease mechanism and predicting most influential genes as potential drug targets or biomarkers of the disease condition of interest. As an exemplary analysis, we performed an application of the software over LNCaP datasets and, out of approximately 25000 genes, predicted CXCR7 and CXCR4 together as drug targets of LNCaP prostate cancer dataset. We further successfully validated them with our initial wet-lab experiments. The introduced software can be applied to all the diseases, especially cancer, with gene expression data of two different conditions (e.g. tumor vs normal) and thus has the potential of a global benefit. As a distinct remark, our software provide the causal disease mechanism with multiple potential drug-targets rather than a single independent target prediction.

**Availability:** The introduced R software package for the analysis is available in CRAN at https://cran.r-project.org/web/packages/dc3net and also at https://github.com/altayg/dc3net

## Background

Treatment options for many deadly cancers remained limited and the productivity of existing drug development pipelines has been declining as drug discovery efforts are mostly focusing on previously validated drug target proteins. Therefore, there is an urgent need for the identification of new cancer drug targets (Jeon, 2014). Nowadays, the identification of novel oncogenes or tumor suppressor genes has become popular in tumorigenesis studies in understanding molecular mechanisms that drive disease progression (Ren, 2015). Since the development of improved cancer therapies is often mentioned as an urgent unmet medical need and more modern methodologies such genetic interactions are being therapeutically exploited to identify novel targeted treatments for cancer (Benstead-Hume, 2017). Understanding the working mechanism of molecules in normal cell physiology and pathogenesis allows subtle drug development and helps treatment of a disease, such as cancer (Altay, 2010; Rual, 2005; Schadt, 2009). The advent of systems and network biology enable us to capture interactions occurring within a cell, which can be represented as gene networks. Computational analysis of the networks provides key insights into biological pathways and cellular organization (Altay, 2011).

The biological processes at the gene level are very complex structures as genes dynamically interact with each other. The interactions of these molecules have been changing significantly over time and in different cell conditions such as from normal to cancer (Emmert-Streib, 2012; Califano, 2011). A single gene can participate in different biological processes and regulate different genes at different times. However, diseases are usually consequences of interactions between multiple molecular processes, rather than an abnormality in a single gene (Menche, 2015).

Gene regulatory networks hold the potential to identify specific subnetworks that are dysfunctional in the disease state of a cell. The identification of differences between disease and healthy tissues may provide key insights into the underlying mechanisms of diseases (de la Fuente, 2010; Ideker, 2012). For this purpose, several methods have been proposed in the literature. However, none of these methods, to our knowledge, provided direct physical interactions in the inferred differential network. Differential C3NET (DC3NET) algorithm was introduced to match that need (Altay, 2011), which is based on the very conservative gene regulatory network (GRN) inference algorithm C3NET (Altay, 2010). In order to ease the usage of DC3NET, we developed an R software, dc3net, and introduce it along with its application on a real dataset in this paper. This study is not an introduction of the algorithm, DC3NET, which is already available in its original publication. Rather, in this study, we developed an R software package of the algorithms for wider utilization of the algorithm. We also applied it to a real dataset and performed wet-lab validations for the top two target predictions. The results suggest that, our introduced software might also provide new drug targets for other diseases if applied with our software as presented in this study. This study aims to present the software to researchers who might apply it to their gene expression data with their disease of interest. Therefore, it has the potential of global health benefit as it may cause finding new drug targets or biomarkers for other diseases. As an application example, we applied it to a publically available LNCaP prostate cancer dataset (GSE18684).

Prostate cancer is the second most common cancer in the male population, with an estimated 417,000 new cases diagnosed each year in Europe (Ferlay, 2013). The activation of androgen receptor (AR) through androgens plays a crucial role in the development and progression of prostate cancer (Kaur, 2016; Anantharaman, 2015; Choudhary, 2011; Massie, 2011). For early detection of prostate cancer, prostate-specific antigen (PSA) screening method has been used widely as a diagnostic tool (Karatas, 2015). However, PSA fails to discriminate indolent disease, which results in over-diagnosis and this may lead to poor prognosis (Abou-Ouf, 2015; Ma, 2015; Myers, 2015). Furthermore, there is no evidence showing that the PSA screening reduces the incidence of death and the underlying mechanism of prostate cancer progression remains largely unknown (Cannistraci, 2014;Ren, 2015).

In the present study, differential gene network analysis has been performed to detect the differences between androgen-stimulated and androgen-deprived prostate cancer cells. We have used differential network inference software tool, dc3net, based on the algorithm DC3NET (Altay, 2011) and used it to infer androgen stimulated prostate cancer specific differential network which can be seen in Figure 1. We have performed validation study on our results via pathway analysis and with direct support from literature that substantiate our blind estimation on prostate cancer. Additionally, our wet-lab experiments show that silencing CXCR7 and CXCR4 dramatically reduce the proliferation of LNCaP prostate cancer cells; this supports our computational drug target prediction.

These two predicted targets have been validated by our experiments and also have strong support from the previous literature. This suggests that our analysis and the software we present in this study are useful in identification of potential drug targets or biomarkers. Our approach ideally apply to cases where there is a comparative disease-related gene expression dataset and a reference dataset, such as cancer and healthy cell conditions.

## 2. Materials and methods

### 2.1. Microarray data and data preprocessing

In order to investigate the alterations in androgen stimulated prostate cancer cells compared with androgen-deprived prostate cancer cells, the microarray dataset, GSE18684 generated by Massie et al. (2011), was obtained from the Gene Expression Omnibus (GEO, http://www.ncbi.nlm.nih.gov/geo). The expression profile included 96 samples, comprising 20 androgen-deprived tissue samples and 76 tissue samples with androgen stimulated prostate cancer. Sample size of 76 is sufficient to infer gene network with maximum performance but sample size of 20 may provide reduced performance according to (Altay, 2012).

The raw microarray data was analyzed using R software v.3.2.2 (http://www.r-project.org) and further processed using Affy package in Bioconductor (Gautier, 2004). Background correction, quantile normalization, and probe summarization processes were performed by the Robust Multi-array Average (RMA) algorithm, in order to obtain gene expression matrix for each dataset (normal and tumor) (Irizarry, 2003).

In microarray technology, multiple probes can represent a single gene. In theory and mostly in practice, those kinds of probes have highest association scores among them, which can cause errors in the inference algorithm *c3net* since it infers only the highest correlated pair for each significant gene. When one works at the probe level data, one should address this issue. In order to eliminate this problem, we filter the association matrix by setting zero for the mutual information score for those probe pairs that correspond to the same gene (Altay, 2010). We have developed a function in the package to practically overcome this issue.

### 2.2. Differential network analysis

In order to perform genome-wide differential gene network analysis, we used the software tool dc3net, which is available in CRAN (https://cran.r-project.org/web/packages/dc3net). Briefly, the dc3net algorithm takes two different microarray gene expression data sets as input (e.g. one as tumor and the other as control). Then, two different gene networks are inferred by applying C3NET (Altay, 2010, Altay, 2010) to each dataset. In order to compute dependency scores among genes, we used PBG that computes mutual information values that provide sufficient performance with very low complexity (Kurt, 2014).

In the final step, these two networks are compared and tumor differential network and common network are inferred. The tumor differential network, *difnet*, is inferred by selecting only the interactions of tumor differential network that does not strong association scores in the control network. The common network, *comnet*, is determined by selecting the overlapping or closer interactions in value or rank between the two networks (Altay, 2011).

In this study, we used the differential network inference all-in-one function *dc3net* with those parameters as follows, where the further details can be seen in the CRAN repository:

*“dc3net(test_data, control_data, probe_names, gene_names, method="cutoff", methodValue=0, itNum=1, rankDif=2000, percentDif=0.3, rankdCom=100, percentCom=0.6, probFiltered=FALSE, visualization=TRUE)”*

### 2.3. Gene ontology analysis

Gene Ontology (GO) enrichment analysis based on Gene Ontology database (http://www.geneontology.org) was performed to investigate the biological roles of the genes in the differential network (da Huang, 2009). To further assess the signaling pathway of the genes, we subsequently performed Kyoto Encyclopedia of Genes and Genomes (KEGG, http://www.genome.jp/kegg) pathway enrichment analysis. The two analysis were performed using The Database for Annotation, Visualization and Integrated Discovery (DAVID, https://david.ncifcrf.gov) which is a powerful bioinformatics tool to find out functions of interested genes (Dennis, 2003). The enrichment analyses required >5 genes to be present and p<0.05 for a term to be considered significant.

**Figure 1.**
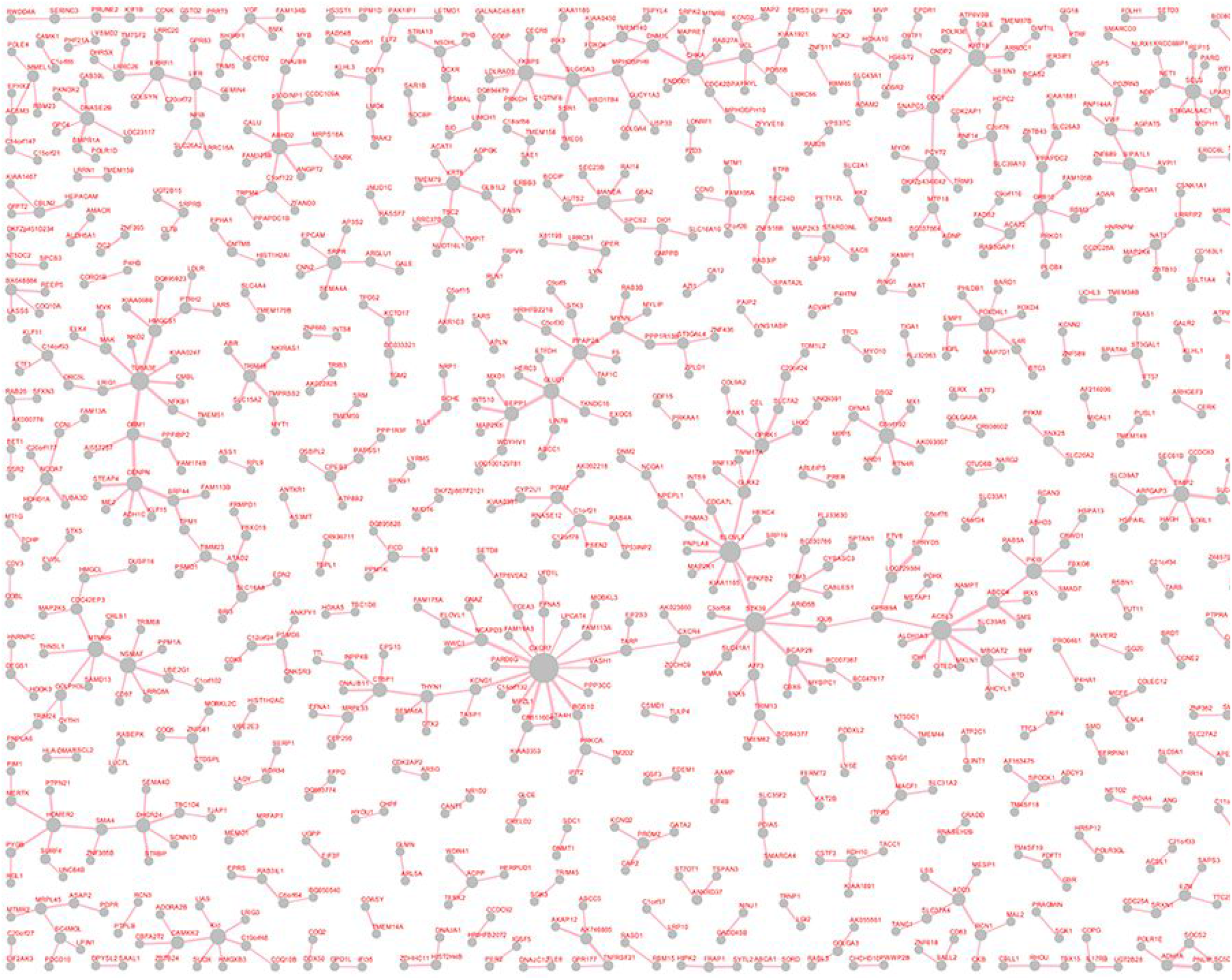
Genome-wide androgen-stimulated prostate-specific differential network with 891 interactions

## 3. Results

### 3.1. Inferring differential network

The androgen-stimulated prostate cancer differential gene network with 891 interactions was inferred using the dc3net tool where details of its usage can be seen in the Supplementary File. The largest independent subnetwork with 119 interactions was also extracted from the differential network and plotted as in Figure 3.

### 3.2. Enrichment analysis

#### 3.2.1. Functional enrichment analysis

To investigate the functions of the genes in the androgen-stimulated prostate cancer differential gene network, GO and KEGG pathway analyses were performed. A total 184 terms were retrieved from the DAVID online analytical tool.

The top ten GO terms ranked by statistical significance were listed in Table 1. GO analysis revealed that genes associated with sterol biosynthetic process (GO:0016126; p=5.05 e-08), protein transport (GO:0015031; p=2.57 e-07) and establishment of protein localization (GO:0045184; p=3.80 e-07) were significantly enriched top three GO terms among biological processes, while for molecular functions, nucleotide binding (GO:0000166; p=7.08 e-05), purine nucleotide binding (GO:0017076; p=5.22 e-04) and purine ribonucleotide binding (GO:0032555; p=9.11 e-04) were significantly enriched, and with regards to cellular components, genes associated with endoplasmic reticulum (GO:0005783; p=2.66 e-13), endoplasmic reticulum part (GO:0044432; p=1.16 e-06) and organelle membrane (GO:0031090; p=1.48 e-04) were significantly enriched (Table 1, Fig. 2A). In Figure 2, we present the functional annotation of the significantly enriched genes in the differential network seen in Figure 1.

**Table 1.** GO terms of differential network (top 10).

| GO ID | GO term | No. of genes | p |
| --- | --- | --- | --- |
| Biological processes |  |  |  |
| GO:0016126 | sterol biosynthetic process | 14 | 5.05E-08 |
| GO:0015031 | protein transport | 82 | 2.57E-07 |
| GO:0045184 | establishment of protein localization | 82 | 3.80E-07 |
| GO:0046907 | intracellular transport | 73 | 3.96E-07 |
| GO:0006695 | cholesterol biosynthetic process | 11 | 1.25E-06 |
| GO:0016125 | sterol metabolic process | 21 | 1.87E-06 |
| GO:0006886 | intracellular protein transport | 47 | 2.61E-06 |
| GO:0008104 | protein localization | 87 | 4.16E-06 |
| GO:0034613 | cellular protein localization | 49 | 6.76E-06 |
| GO:0008203 | cholesterol metabolic process | 19 | 7.26E-06 |
| Cellular components |  |  |  |
| GO:0005783 | endoplasmic reticulum | 117 | 2.66E-13 |
| GO:0044432 | endoplasmic reticulum part | 46 | 1.16E-06 |
| GO:0031090 | organelle membrane | 97 | 1.48E-04 |
| GO:0005789 | endoplasmic reticulum membrane | 33 | 2.11E-04 |
| GO:0042175 | nuclear envelope-endoplasmic reticulum network | 34 | 2.63E-04 |
| GO:0005739 | mitochondrion | 92 | 9.73E-04 |
| GO:0005829 | cytosol | 109 | 9.89E-04 |
| GO:0005792 | microsome | 28 | 1.24E-03 |
| GO:0005624 | membrane fraction | 71 | 1.61E-03 |
| GO:0042598 | vesicular fraction | 28 | 1.90E-03 |
| Molecular Function |  |  |  |
| GO:0000166 | nucleotide binding | 174 | 7.08E-05 |
| GO:0017076 | purine nucleotide binding | 147 | 5.22E-04 |
| GO:0032555 | purine ribonucleotide binding | 140 | 9.11E-04 |
| GO:0032553 | ribonucleotide binding | 140 | 9.11E-04 |
| GO:0000287 | magnesium ion binding | 42 | 4.10E-03 |
| GO:0001883 | purine nucleoside binding | 119 | 5.75E-03 |
| GO:0003924 | GTPase activity | 23 | 6.81E-03 |
| GO:0001882 | nucleoside binding | 119 | 7.19E-03 |
| GO:0005524 | ATP binding | 110 | 7.74E-03 |
| GO:0004674 | protein serine/threonine kinase activity | 39 | 8.48E-03 |
| FDR: false discovery rate; GO: gene ontology. |  |  |  |

**Figure 2.**
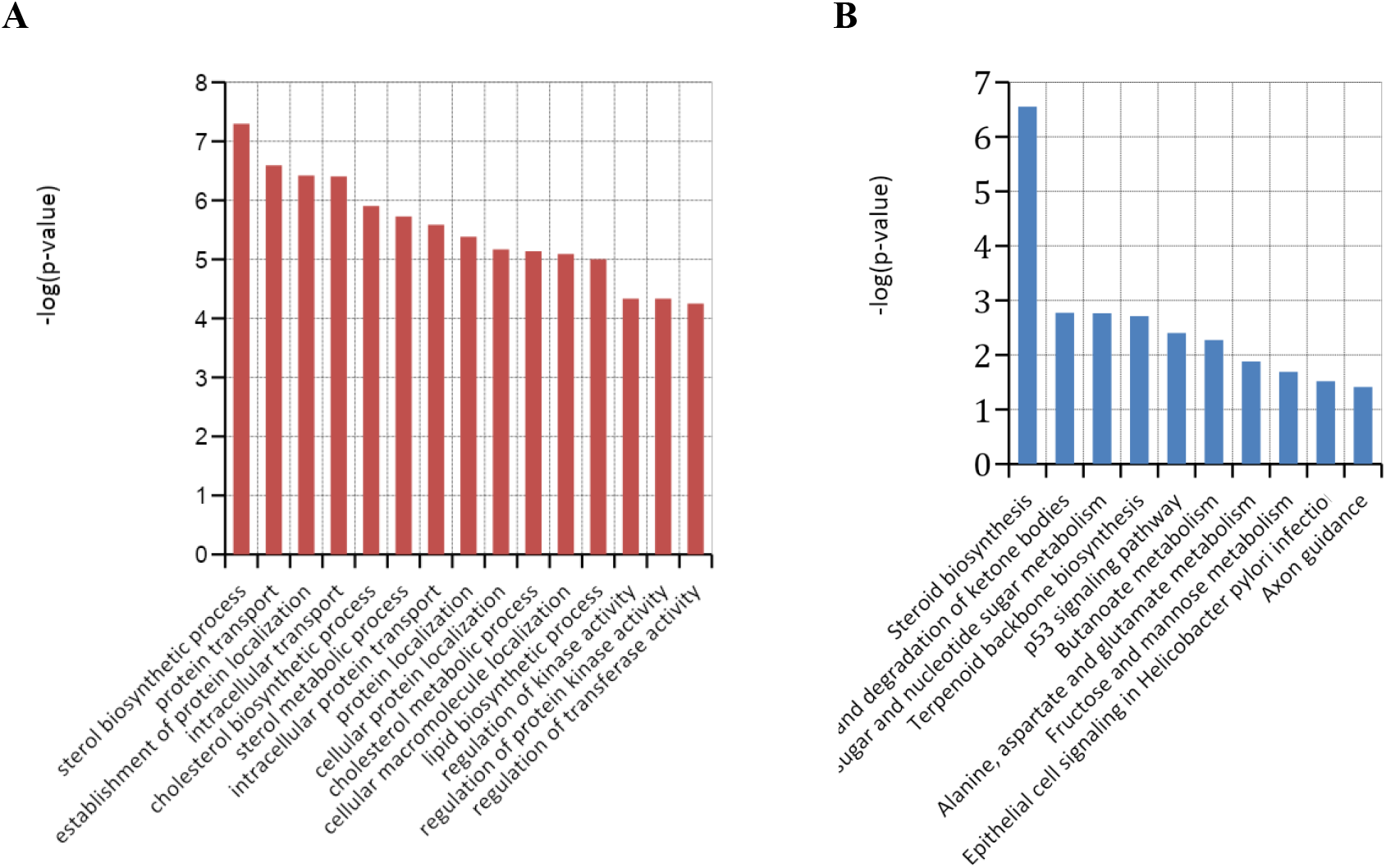
Functional annotation of significantly enriched genes in differential network. (A) The top 10 enriched gene ontology categories for biological processes; (B) The enriched Kyoto Encyclopedia of Genes and Genomes pathways.

Next, the genes found in the androgen-stimulated prostate cancer differential gene network were submitted to DAVID server to identify significantly enriched KEGG pathways (Kanehisa, 2000; Kanehisa, 2012). The KEGG pathways that were found significantly enriched (p<0.05) are shown in Table 2. Pathway analysis revealed that the genes in the androgen stimulated prostate cancer difnet were significantly enriched in ten terms. The most significant three terms were those involved in steroid biosynthesis (p=2.80 e-07), synthesis and degradation of ketone bodies (p=1.646523 e-03), and amino sugar and nucleotide sugar metabolism (p=1.73 e-03) processes (Fig. 2B).

**Table 2.** Significant KEGG pathways in the androgen stimulated prostate cancer specific differential network.

| KEGG ID | KEGG term | No. of genes | p | Genes |
| --- | --- | --- | --- | --- |
| hsa00100 | Steroid biosynthesis | 10 | 2.80E-07 | TM7SF2, CEL, EBP, SQLE, LSS, SC5DL, DHCR24, FDFT1, SC4MOL, NSDHL |
| hsa00072 | Synthesis and degradation of ketone bodies | 5 | 1.69E-03 | HMGCS2, HMGCS1, ACAT2, ACAT1, HMGCL |
| hsa00520 | Amino sugar and nucleotide sugar metabolism | 10 | 1.71E-03 | PGM2, GMPPB, GALK1, PGM3, GNPDA1, CMAS, GFPT1, GFPT2, HK2, GALE |
| hsa00900 | Terpenoid backbone biosynthesis | 6 | 1.95E-03 | HMGCS2, HMGCS1, FDPS, MVK, ACAT2, ACAT1 |
| hsa04115 | p53 signaling pathway | 12 | 3.94E-03 | CCNE2, BID, PPM1D, TSC2, SIAH1, CDK6, CCNG1, GADD45B, THBS1, IGFBP3, GADD45A, SESN3 |
| hsa00650 | Butanoate metabolism | 8 | 5.33E-03 | ACSM3, HMGCS2, ALDH5A1, HMGCS1, ABAT, ACAT2, ACAT1, HMGCL |
| hsa00250 | Alanine, aspartate and glutamate metabolism | 7 | 1.32E-02 | ASS1, ALDH5A1, GFPT1, GLUD1, GFPT2, ABAT, ALDH4A1 |
| hsa00051 | Fructose and mannose metabolism | 7 | 2.05E-02 | MTMR2, GMPPB, SORD, PFKFB2, HK2, PFKM, MTMR6 |
| hsa05120 | Epithelial cell signaling in Helicobacter pylori infection | 10 | 3.03E-02 | IGSF5, ATP6V0E1, LYN, ATP6V1E1, MAP2K4, NFKB1, PAK1, JAM3, ATP6V0A2, ATP6V0B |
| hsa04360 | Axon guidance | 15 | 3.89E-02 | NRP1, EFNB3, EFNA1, EFNB2, DPYSL2, EPHA1, SEMA6A, NCK2, PAK2, CXCR4, PPP3CC, EFNA5, PAK1, SEMA4D, SEMA4A |

KEGG: Kyoto Encyclopedia of genes and genomes

In order to further evaluate the biological roles of the genes in the independent subnetworks of the genome-wide androgen-stimulated prostate cancer difnet, we performed KEGG analysis for the largest subnetwork. As shown on Figure 3, this subnetwork comprises 119 interactions with CXCR7, STK39, ELOVL3 and ACSL3 at the center of the largest hubs. KEGG analysis of the genes included in the subnetwork revealed a highly significant association with axon guidance pathway (p=1.71 e-03), which was also found significantly enriched in the whole differential network. Furthermore, pathways involved in Fc gamma R-mediated phagocytosis (p=2.69 e-02) and Endocytosis (p=3.62 e-02) were also highly enriched (Table 3). Interestingly, these two pathways were not found significantly enriched in the whole differential network.

**Figure 3.**
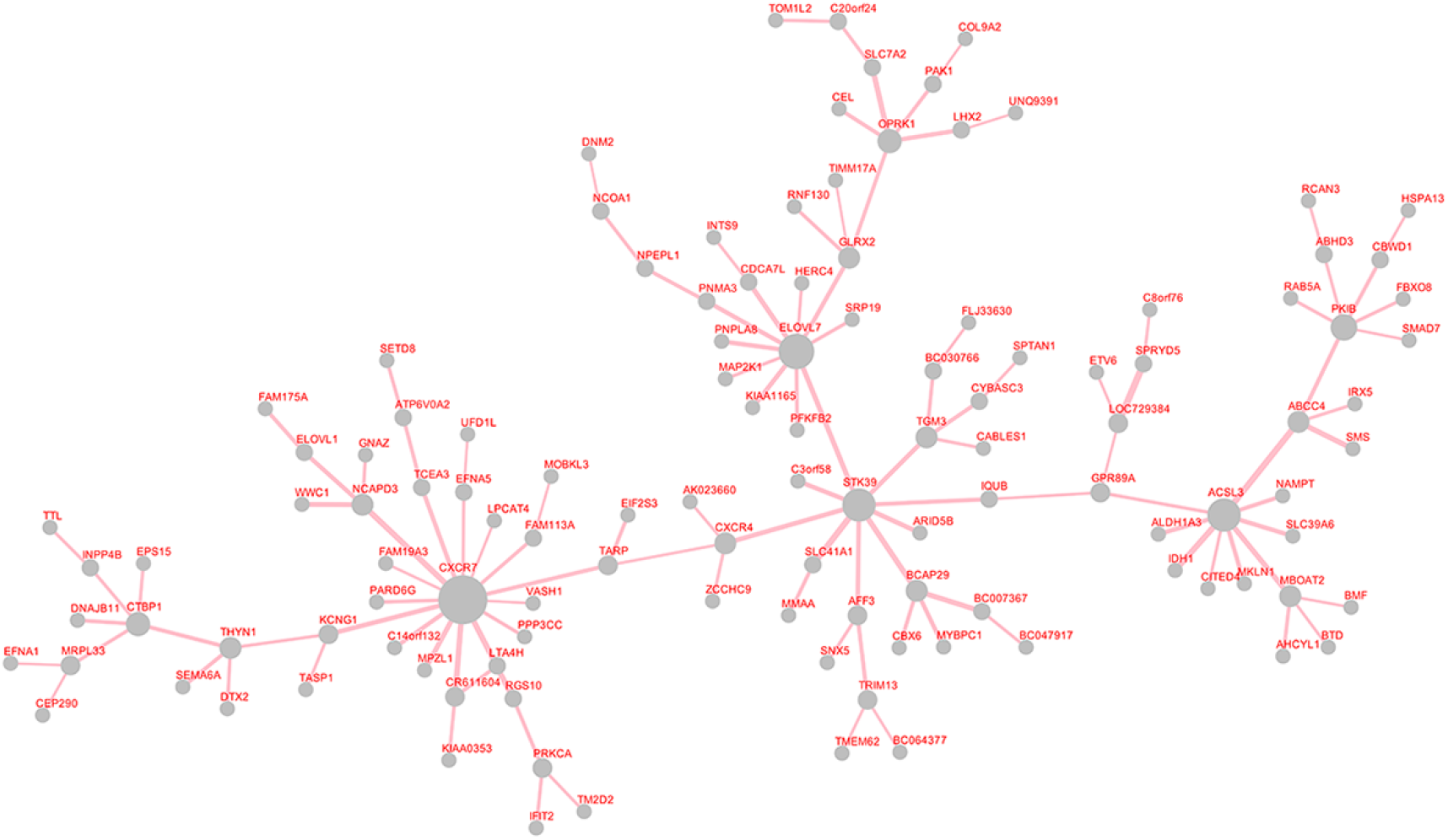
The largest connected subnetwork of the androgen-stimulated prostate cancer difnet. This subnetwork might have an important role in human prostate cancer as being the largest connected subnetwork with 119 edges in tumor difnet.

**Table 3.** Significant KEGG pathways in the largest subnetwork of the androgen stimulated prostate cancer specific differential network.

| KEGG ID | KEGG term | No. of genes | p | Genes |
| --- | --- | --- | --- | --- |
| hsa04360 | Axon guidance | 6 | 1.71E-03 | SEMA6A, CXCR4, EFNA1, PPP3CC, EFNA5, PAK1 |
| hsa04666 | Fc gamma R-mediated phagocytosis | 4 | 2.69E-02 | PRKCA, MAP2K1, PAK1, DNM2 |
| hsa04144 | Endocytosis | 5 | 3.62E-02 | EPS15, CXCR4, RAB5A, PARD6G, DNM2 |

KEGG: Kyoto Encyclopedia of genes and genomes

In order to investigate biological functions of the genes in the differential network, GO and KEGG pathway enrichment analyses were performed. The pathway analysis is important since it improves disease classification and reveals novel insights about a disease state (Myers, 2015). The results showed that sterol biosynthetic process was the most significantly enriched GO term for biological process. To further evaluate the biological roles of the genes in the differential network, KEGG pathway analysis was performed. According to the KEGG analysis, steroid biosynthesis was the most significant pathway (p=2.80 e-07). It contains ten genes in our network: TM7SF2, CEL, EBP, SQLE, LSS, SC5DL, DHCR24, FDFT1, SC4MOL and NSDHL. The relation of steroid biosynthesis and prostate cancer is reported in many studies. The ligand activation of the androgen receptor plays an important role in the progress of castration-resistant prostate cancers. The similarities and differences in glandular androgen synthesis provide direction for the development of new treatments (Migita, 2009; Sharifi, 2012; Auchus, 2012; Ferraldeschi, 2013).

Of second highest significance was the pathway for synthesis and degradation of ketone bodies (p=1.63 e-03), which contains five genes: HMGCS2, HMGCS1, ACAT2, ACAT1 and HMGCL. In the study conducted by Lin et. al. (2005), this pathway was found up-regulated in androgen-independent CL1 cells (model for late-stage prostate cancer) when compared to androgen-dependent LnCaP (model for early-stage prostate cancer) cells.

We have also examined the other significant pathways, and found that Amino sugar and nucleotide sugar metabolism (Priolo, 2014), p53 signaling pathway (Chappell, 2012; Gupta, 2012; Stegh, 2012), Butanoate metabolism (Stoss, 2008; Romanuik, 2010), Alanine, aspartate and glutamate metabolism (Priolo, 2014), and Axon guidance (Choi, 2014) pathways were shown to be associated with the prognosis of prostate cancer. In these pathways, the p53 signaling pathway plays a critical role in cancer’s response to chemotherapy and tumor growth. Inactivation of the tumor suppressor gene p53 is widely observed in more than 50% of human cancers including prostate cancer. The disruption of the p53 signaling pathway is one of the vital turning point for the survival of advanced prostate cancer cells during therapies. By enabling DNA repair, it was observed that p53 blocks cancer progression by provoking transient or permanent growth arrest (Chappell, 2012; Gupta, 2012; Stegh, 2012). However, three pathways, Terpenoid backbone biosynthesis (hsa00900), Fructose and mannose metabolism (hsa00051) and Epithelial cell signaling in Helicobacter pylori infection (hsa05120), have not previously been related to prostate cancer.

KEGG analysis for the largest independent subnetwork revealed much more interesting results that may show that it has the most important role in the prostate cancer. Axon guidance (hsa04360) pathway, which has also been found to be significantly enriched in the whole differential network, is known to have tumor suppressor genes and therefore related with tumor growth (Mehlen, 2011). Axon guidance molecules are validated as tumor suppressor in the breast cancer and show promise as breast cancer diagnostic markers as well as potential therapeutic targets (Mehlen, 2011; Harburg, 2011). In the study conducted by Choi, axon guidance pathway was shown to be involved in prostate cancer tumorigenesis (Choi, 2014). In addition, Savli et al. reported that axon guidance signaling pathway was the most significant down-regulated canonical pathway in prostate cancer (Savli, 2008). The second significantly enriched pathway was Fc gamma R-mediated phagocytosis (hsa04666). This pathway was found as the highest significant pathway in prostate cancer and have been referred as being involved in the pathological development of prostate cancer (Jia, 2012). In the literature, the endocytosis (hsa04144) pathway was also found as related to prostate cancer. The importance of understanding the regulation between signal transduction and endocytosis pathways, and also how the breakdown of this integrated regulation contributes to cancer development was emphasized (Bonaccorsi, 2007).

In brief, our top four strongest hubs with blind predictions have strong support from the literature. In the future, one can further investigate these top 4 hubs together in LNCaP cells, to elucidate their role in prostate cancer biology. Additionally, KEGG pathway analysis on our androgen-stimulated prostate cancer-specific differential network has revealed outstanding results. According to the KEGG pathway analysis results, out of ten most significantly enriched pathways, seven are already known to have strong association with prostate cancer. We performed disease-specific network inference with a blind prediction, and it strongly correlated with the pathway analysis. We suggest that the three unrelated pathways in prostate cancer deserve further experimental investigation, in order to reveal their relation to prostate cancer biology. More interestingly, KEGG pathway analysis in the largest independent subnetwork of our androgen-stimulated prostate cancer difnet, has revealed three pathways where all of them are known to have strong association with prostate cancer. This shows the highly accurate performance of the differential network analysis tool dc3net. We concluded that this is the largest subnetwork is one of the key mechanisms in our androgen-stimulated prostate cancer difnet. Our experimental results on the top two targets provide experimental evidence that supports our predictions.

#### 3.2.2. Literature search of the top hub targets

Top four hub nodes identified in the present study, have been strongly associated with prostate cancer metastatic process, including CXCR7, STK39, ELOVL7 and ACSL3. Hub nodes are genes that are highly connected with other genes and they were proposed to have important roles in biological development. Since hub nodes have more complex interactions than other genes, they may have crucial roles in the underlying mechanisms of disease (Guo, 2015). Identification of hub genes involved in progression of prostate cancer may lead to the development of better diagnostic methods and providing therapeutic approaches.

According to our analysis, CXCR7 (chemokine (C-X-C motif) receptor 7) is by far the top hub gene in the androgen-stimulated differential network and it is also part of the largest independent subnetwork as seen in Figure 3. In (Wang, 2008), it is reported that staining of high-density tissue microarrays showed increased levels of CXCR7/RDC1 expression in correlation to tumor aggressiveness. Also, *in vitro* and *in vivo* studies with prostate cancer cell lines propose that alterations in CXCR7/RDC1 expressions are associated with enhanced invasive and adhesive activities in addition to a survival advantage. Along other papers on CXCR7 (Zheng, 2010), it was shown that increased CXCR7 expression was found in hepatocellular carcinoma (HCC) tissues. Knockdown of CXCR7 expression with shRNA significantly inhibited SMMC-7721 angiogenesis, adhesion and cells invasion and also down-regulation of CXCR7 expression led to reduction of tumor growth in a xenograft model of HCC (Zheng, 2010). Another study demonstrated that the IL-8–regulated CXCR7 stimulated EGFR Signaling to promote prostate cancer growth (Singh, 2011). In a study conducted by Yun *et al*., it is reported that CXCR7 expression was increased in most of the tumor cells compared with the normal cells and is involved in cell proliferation, migration, survival, invasion and angiogenesis during the initiation and progression of many cancer types including prostate cancer (Yun, 2015). It was also shown in (Lin, 2014) that elevated expression of CXCR7 contributes to HCC growth and invasiveness via activation of MAPK and angiogenesis signaling pathways and they also conclude that targeting CXCR7 may prevent metastasis and provide a potential therapeutic strategy for hepatocellular carcinoma (HCC).

It was also observed that CXCR7/RDC1 levels are regulated by CXCR4 (Singh, 2011). This is a very interesting supporting information from literature for our blind estimation because in our predicted largest independent subnetwork, as shown in Figure 3, CXCR7 and CXCR4 appear to be very close and interacting over only one gene. Although CRCX4 is not a hub gene, it appears to be a bridge that connects both halves of the largest subnetwork. According to KEGG analysis, CXCR4 was found in the gene list of two different significantly enriched KEGG pathways, axon guidance and endocytosis, which are strongly associated with prostate cancer (Table 3). It is also reported (Shanmugam, 2011) that inhibition of CXCR4/CXCL12 signaling axis by ursolic acid leads to suppression of metastasis in transgenic adenocarcinoma of mouse prostate model and CXCR4 induced a more aggressive phenotype in prostate cancer (Miki, 2007). In another study, it is reported that CXCR4 and CXCR7 have critical roles on mediating tumor metastasis in various types of cancers as both being a receptor for an important *α*-chemokine, CXCL12 (Sun, 2010). Furthermore, a more recent study concluded that CXCR4 plays a crucial role in cancer proliferation, dissemination and invasion and the inhibition of CXCR4 strongly affects prostate cancer metastatic disease (Gravina, 2015). The strong literature that shows the important role of CXCR4 in cancer lead to develop drugs that target it. For example, X4 Pharmaceuticals is a clinical stage biotechnology company developing novel CXCR4 inhibitor drugs to improve immune cell trafficking to treat cancer and rare diseases. The company described that CXCR4 protein “acts as a beacon to attract cells to surround a tumor, effectively hiding the tumor from the body’s T cells that would otherwise destroy them” and. also mentioned that X4 company is beginning human trials using CXCR4 inhibitors which aims to develop a therapy to block the protein, CXCR4 (https://www.biospace.com/article/x4-pharma-uses-37-5-million-to-push-cancer-therapies-into-human-trials-/, 2015).

Although there is strong support for the top two predicted targets, they are part of large subnetwork that might well be the gene interaction mechanism that drives the tumor condition. Therefore, we also searched the literature for the other hubs as the next potential targets and found considerable support for them too.

The second most likely prediction as hub was STK39 (serine threonine kinase 39). Among others, in (Hendriksen, 2006) it is reported that lower mRNA expression of STK39 in primary prostate tumors was correlated with a higher incidence of metastases after radical prostatectomy. In (Balatoni, 2009), it is stated that STK39 encoded protein SPAK, regulates cell stress responses, and microarray studies identified reduced SPAK expression in treatment-resistant breast cancers and metastatic prostate cancers, suggesting that its loss may play a role in cancer progression. They showed that epigenetic silencing of STK39 in B-cell lymphoma inhibits apoptosis from genotoxic stress in cancer. STK39 is also identified as hypertension susceptibility gene (Wang, 2008).

ELOVL7 (fatty acid elongase 7) was reported that it could be involved in prostate cancer cell growth and survival through the metabolism of SVLFAs and their derivatives, could be a key molecule to elucidate the association between fat dietary intake and prostate carcinogenesis, and could also be a promising molecular target for development of new therapeutic or preventive strategies for prostate cancers (Tamura, 2009).

ACSL3 (acyl-CoA synthetase long-chain family member 3) was reported to be one of the androgen-regulated genes and it is shown that ACSL3 is slightly up-regulated in primary prostate tumors and strongly repressed in metastatic cancer (Marques, 2011). It also states that ACSL3, ELOVL5 and GLUD1 play a role in the production of prostatic fluid and in secretory function of the prostate. From this literature information, it worth mentioning that we blindly predicted ACSL3, ELOVL7 and GLUD1 as in top eight tumor-specific hubs, which may suggest their collaborative role in this disease from this biological process. There is also a patent that reports that the fusion genes ACSL3 and ETV1 and their expression products can be used as prognostic and diagnostic markers for prostate cancer and as clinical targets for the treatment of prostate cancer (Attard, 2008).

### 3.3. Experimental Validation

As seen in Figure 3, the largest differential subnetwork comprises 119 interactions where CXCR7 is the largest hub gene that is connected to the other three hub genes STK39, ELOVL3 and ACSL3 via the bridge gene CXCR4. Therefore, we predicted CXCR7 and CXCR4 as the most important driving genes of LNCaP with Androgen stimulation. These proteins might be potential drug targets or biomarkers of this cell condition. In order to test this prediction, we performed cell proliferation experiments to see whether silencing of any of these two targets or their combination can reduce cell proliferation. In fact, our wet-lab experimental results show that they are dramatically effective in reducing cell growth of LNCaP prostate cancer cells.

#### 3.3.1 Experimental Methods

LNCaP prostate cancer cells were purchased from ATCC, and were grown in RPMI medium supplemented with 10% charcoal dextran-stripped FBS. Cells were separated into two treatment groups – 1) 2 nM androgen (Sigma Aldrich, cat no R1881) and 2) 0.2% ethanol (vehicle control). After 72h of treatment, cells were seeded into 6-well plate and each treatment group was transfected with siRNA (Origene, cat no SR322283 and SR311311) as follows: A) 50 nM siCXCR7, B) 50 nM siCXCR4, C) 50 nM siCXCR7 + 50 nM siCXCR4, and D) 50 nM siScrambled. Transfection reagent RNAiMAX (Thermo Fisher Scientific, cat no 13778030) was used for transfection according to manufacturer’s instructions. Transfected cells of each treatment group were seeded in triplicates in 96-well plate (500 cells/well) in above-mentioned media. Androgen treatment continued after the transfection. In the course of 7 days, cell proliferation was measured using Alamar blue at wavelength of 530/560 nm. Cell growth of each treatment was normalized to own day 0 (starting point) and converted to percentage growth. GraphPad Prism 5 was used to plot the data.

#### 3.3.2 Experimental Results

LNCaP cells also can grow with or without androgen dependency. Silencing of CXCR7 and CXCR4 demonstrated their effect on proliferation of LNCaP cells in androgen-dependent and androgen-independent state; the effect becomes apparent after day 5 (See Figure 4 and Figure 5). In the androgen-independent state, single silencing of CXCR7 or CXCR4 equally delays cell proliferation by 3-fold (day 7). Double silencing of CXCR7 and CXCR4 further reduced cell proliferation by 2-fold, demonstrating the additive effect (See Figure 5). In the androgen-dependent state, single silencing of CXCR7 reduces cell proliferation by 3-fold, whereas silencing of CXCR4 reduces cell proliferation by more than 4-fold. Double silencing of CXCR7 and CXCR4 does not seem to further reduce cell proliferations, probably because the limit is already reached (See Figure 4). These findings confirm that both CXCR7 and CXCR4 are critical to growth of LNCaP cells with and without androgen dependency. Additionally, the results suggest that although both genes have significant role in LNCaP cell growth, there is difference in mechanism of interacting genes. Indeed silencing of CXCRs seems to cause cell cycle arrest in various prostate cancer cells, as found in an independent study (Singh, 2011). Even though expression patterns of CXCR7 and CXCR4 may vary across various cell types, their silencing unequivocally blocks cell proliferation and tumor growth. Importantly double silencing of CXCR7 and CXCR4 has additive effect on cell growth, an important aspect that draws parallels with our findings.

As seen in Figure 4 and Figure, CXCR7 and CXCR4, our top two predicted targets, have significant effect on growth of LNCaP cancer cells in both androgen-stimulated and deprived states. However, we also observe in Figure 5 that, under the Androgen-deprived condition, the combined silencing of both targets has further additive effect. On the other hand, in the Androgen-stimulated case, silencing of CXCR4 alone provides the combination effect despite the fact that CXCR7 is still very effective. This suggest that there is different gene level interaction mechanism between both conditions though the main driving mechanism as suggested in Figure 1 might be similar in both conditions. The observed difference in the androgen-stimulated condition, the effect of double silencing of CXCR7 and CXCR4 is the same as the single silencing of CXCR4 alone, suggest a serial interaction mechanism as predicted in Figure 3 between CXCR7 and CXCR4. This difference also validates the introduced software, dc3net, that infers condition specific interactions. Differential gene network analysis might have helped to filter out the common interactions of LNCaP and normal cell conditions, which might exist both in cancerous and normal cell conditions.

**Figure 4.**
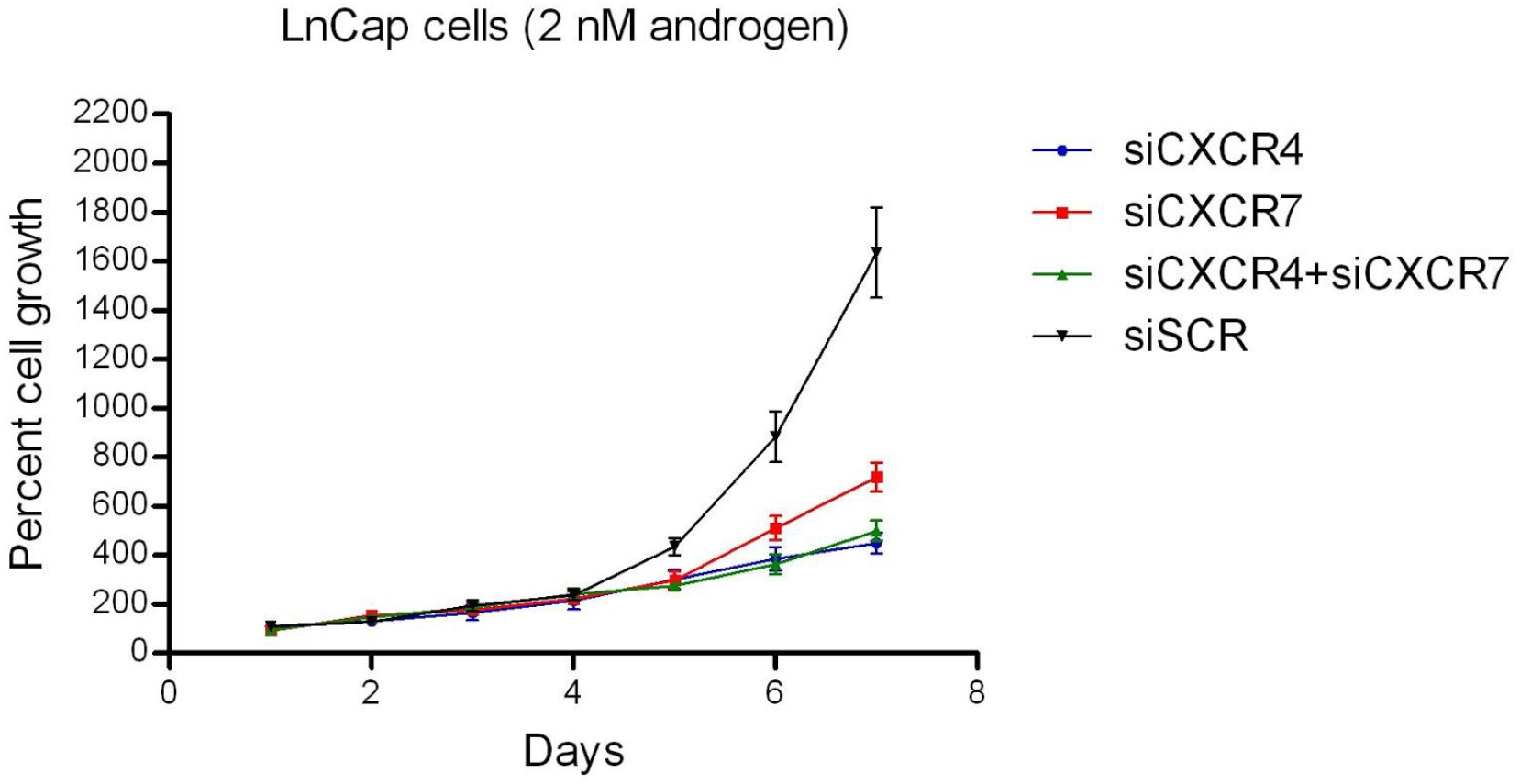
LNCaP growth with Androgen treatment. Effect of CXCR7 and CXCR4 depletion on proliferation of LNCaP prostate cancer cells with androgen treatment. Cells were grown and transfected as described in the Methods section. Plots represent cell growth over the course of 7 days. Each time point is an average of three replicates.

**Figure 5.**
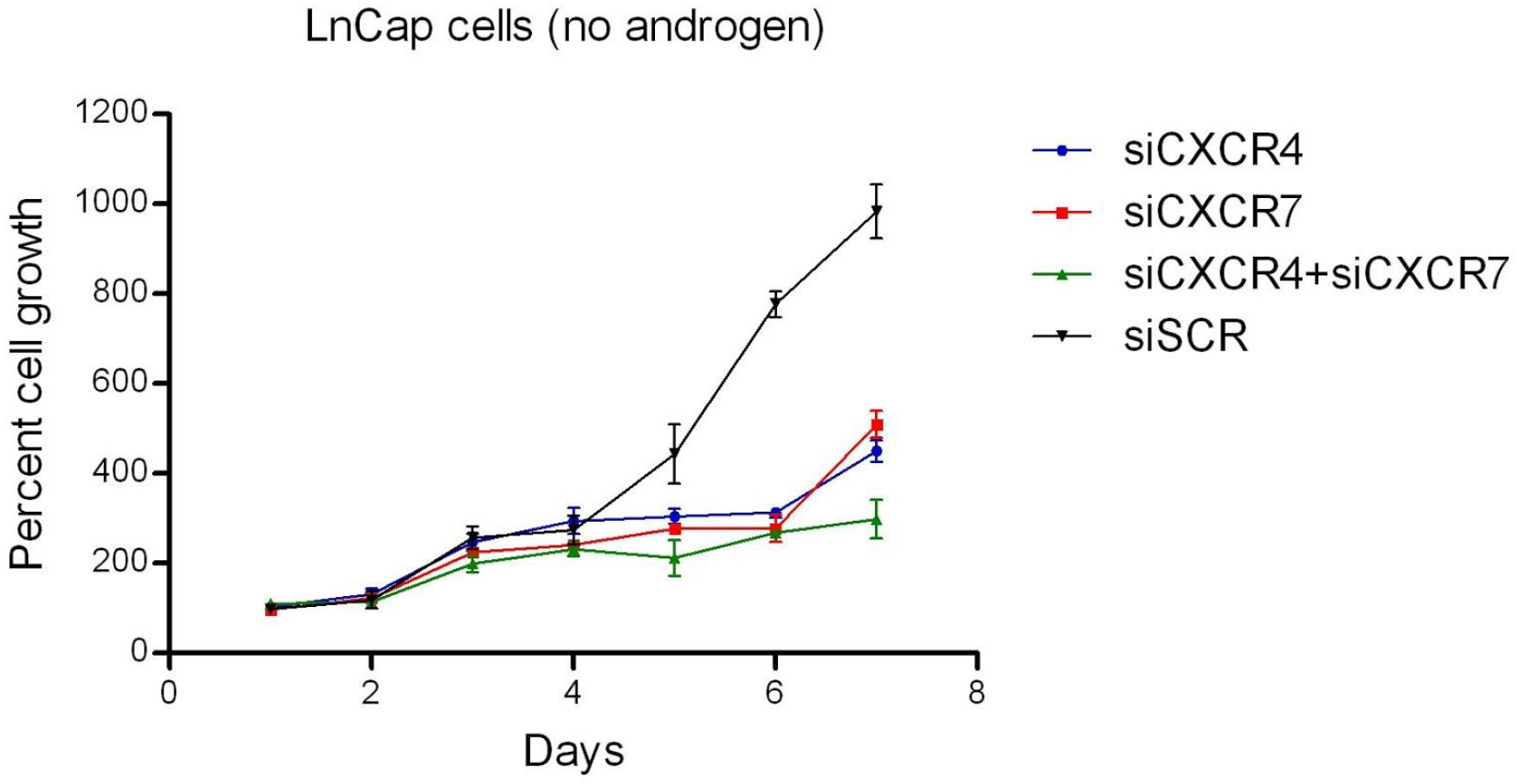
LNCaP growth without Androgen treatment. Effect of CXCR7 and CXCR4 depletion on proliferation of LNCaP prostate cancer cells without androgen treatment. Cells were grown and transfected as described in the Methods section. Plots represent cell growth over the course of 7 days. Each time point is an average of three replicates.

## 4. Discussion

The main goal of this paper is to introduce a software that performs differential network analysis and predicts potential targets that drive the cell condition of interest. In the case of cancer, it might help predicting potential biomarkers or drug targets. In order to show that the software is useful and works, we implemented it on LNCaP prostate cancer datasets with androgen stimulation and androgen-deprived cell conditions. Along with strong literature support, we also performed further wet-lab experiments and validated our top two predictions. Although, both cell types have cancer cells, they have different conditions that are important for the prostate cancer. When we compare them, the normal gene interactions, which might be available in cancer and normal cells are also expected to cancel out each other. However, we expect to observe a different gene interactions mechanism specific to the cell condition of interest (not necessarily different genes). We observed that difference in Figure 4 with compare to Figure 5, where the combined targets effect was observed with one of the targets, which suggest a serial gene interaction mechanism as we observed in Figure 2. Below, we discuss in further detail on the results.

We predicted potential drug targets CXCR7 and CXCR4 in LNCaP prostate cancer, using differential gene network analysis approach, DC3NET. With this study, we present an R software package, dc3net, which was used to implement the algorithm. Our analysis results reveal CXCR7 and CXCR4 as the most promising top two targets. Our wet-lab experimental analysis validated that silencing of CXCR7 and CXCR4 dramatically reduce growth of LNCaP prostate cancer cell, and thus validate them as key driving genes of the LNCaP prostate cancer.

Hence, our blind prediction reveals CXCR7 and CXCR4 as potential drug targets, and our wet-lab results validates our predictions. Indeed, there is strong support from the recent literature on the importance of these targets. Experimental results of (Saha, 2017) suggest that targeting the CXCR4/CXCR7 axis could lead to novel approaches for offsetting the effects of obesity on prostate cancer progression. CXCR7 and CXCR4 were also elaborated in (Xu, 2016). They mentioned that CXCR7 was involved in modulating tumor microenvironment, tumor cell migration and apoptosis, thus revealing that these complex interactions provided insight into targeting CXCR7 as a potential anticancer therapy. Similar to our experimental observation in Figure 5, in (Han, 2017), suppressing CXCR4 was not enough to obstruct osteosarcoma invasion in bone marrow microenvironment, as CXCR7 was activated to sustain invasion. Therefore, they suggested that inhibiting both CXCR4 and CXCR7 could be a promising strategy in controlling osteosarcoma invasion, which perfectly matches our prediction and observation in LNCaP prostate cancer.

## 5. Conclusions

We introduced a software that helps predicting potential biomarkers or drug targets and showed its effectiveness by an application on LNCaP prostate cancer dataset. The results provide significant insight into the molecular mechanisms associated with development of LNCaP prostate cancer. Furthermore, GO and KEGG pathway enrichment analyses identified numerous pathways that may have a role in the prostate cancer, and these findings may contribute to better understanding of the molecular mechanism of this disease and also disclose potential targets for diagnostic and therapeutic use. Our wet-lab data suggest that CXCR7 and CXCR4 as potential drug targets, which is strongly evidenced in the literature. Some of the other predicted targets, which are parts of the same tumor specific subnetwork, in the androgen-stimulated prostate cancer difnet may also have utility as biomarkers or therapeutic targets for prostate cancer and thus merit further investigation.

In conclusion, the software might well be applied to other diseases as well and has the potential to have a wider impact in exploring novel disease specific targets.

## Acknowledgements

This paper is an extension of the pre-print “Genome-Wide Differential Gene Network Analysis R Software And Its Application In LnCap Prostate Cancer”, bioRxiv, April 24, 2017.

We would like to acknowledge the support of The University of Cambridge, Cancer Research UK and Hutchison Whampoa Limited. Parts of this work were funded by CRUK core grant C14303/A17197 and A19274. Gokmen Altay was funded during his research for his contributions.

There was one additional contributor to this manuscript, who has left the study at the final stages without any notification and failed to respond to our several contact attempts; and has not provided information on his affiliation and thus could not be included.

## Supplementary File

### 1. Introduction

The *dc3net* is an R package that infers direct physical interactions of differential gene networks from gene expression datasets of multiple conditions. This supplementary file exemplifies how to use the *dc3net* package and express detailed information on the several workflows with example data sets. The data sets used in this file are available through the dc3net R-package. In the below figure, we present the DC3NET algorithm as block diagram:

**Fig. 1.**
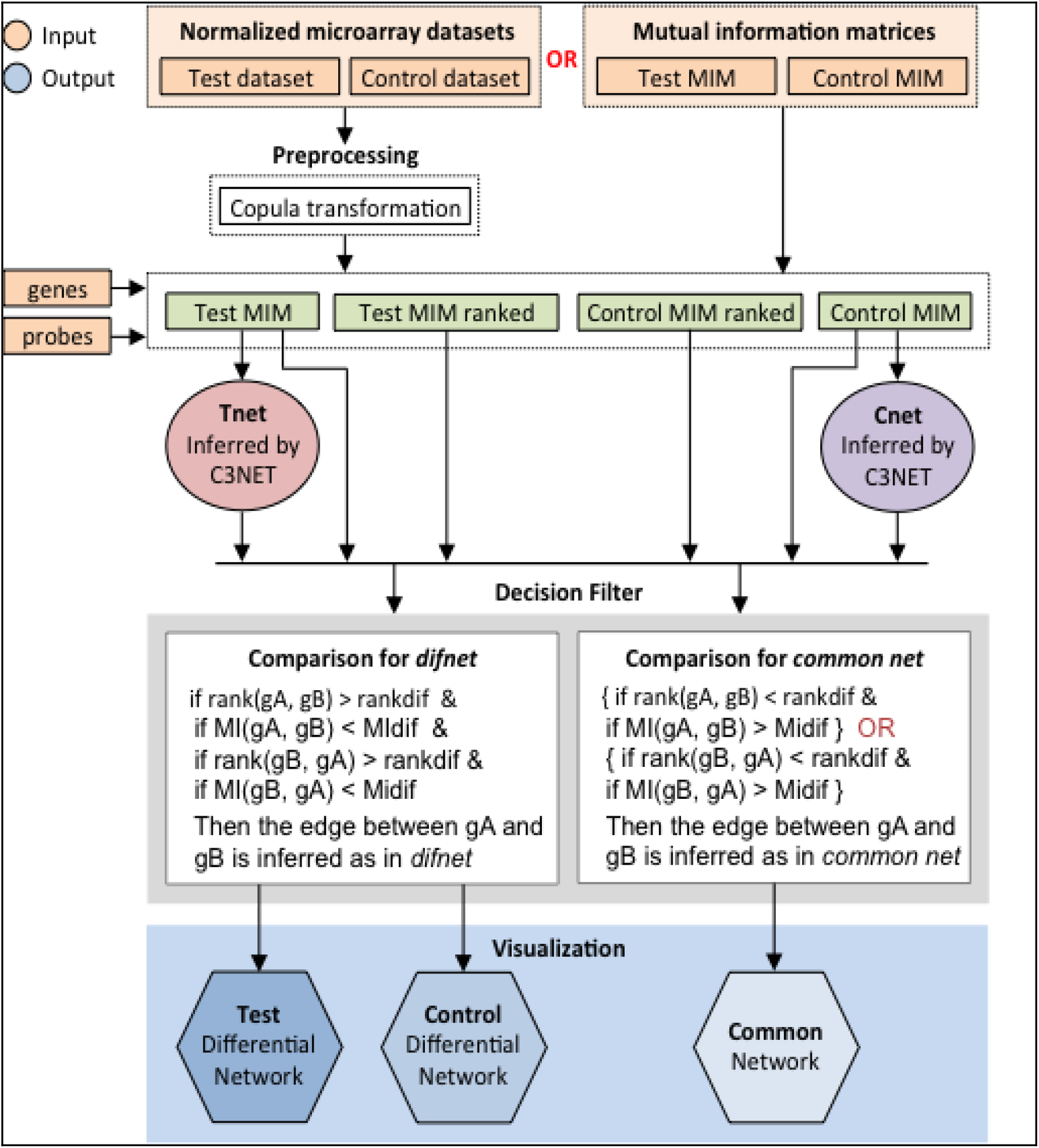
Schematic overview of the dc3net

### 2. Installation of the R package DC3NET

*Dc3net* requires “R 3.2.x and later” and it depends on “*c3net*”, “*igraph*” and “*RedeR*” packages that can be installed from the CRAN and Bioconductor libraries. For the installation of *dc3net*, the user needs to follow some simple installation steps.

1. To download and install dependent packages *c3net, igraph* and *RedeR* from CRAN and Bioconductor (execute in R):

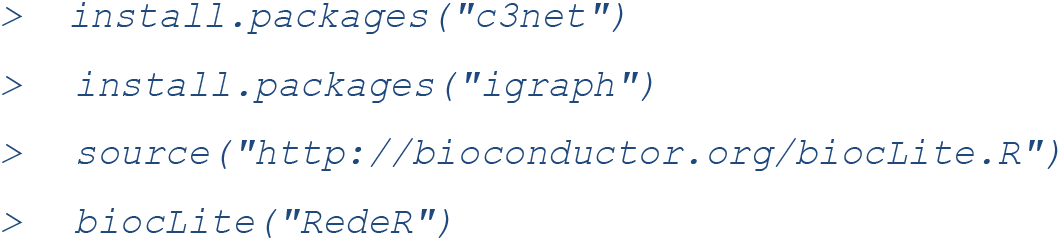
2. Execute the installation command for *dc3net* within R from CRAN or as follows:

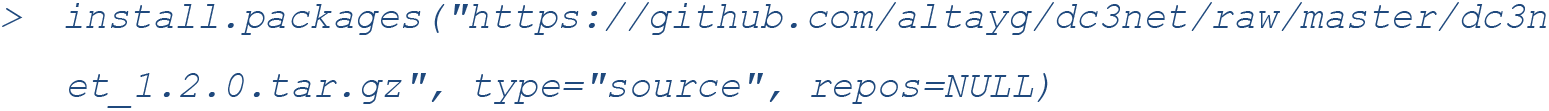
3. For the instructions on the usage of *dc3net*, please check the user manual *dc3net-manual.pdf*
4. To load the library:

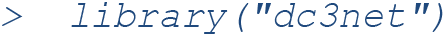

### 3. General guidelines for using DC3NET

A detailed explanation and workflow of DC3NET algorithm can be found in (Altay et al., 2011). Therefore, we do not reproduce the open access text, but we briefly describe the DC3Net algorithm and explain the parameters we used on example data sets.

The required inputs of the package are two different gene expression data sets, probe names and gene names. Users can also use pre-computed test and control mutual information (adjacency) matrices as input. Otherwise, the algorithm takes the two data sets and generates the matrices itself. The data sets need to be normalized together (e.g. using RMA) before using in dc3net. If the input data sets are precomputed mutual information matrices, then the algorithm skip this preprocessing step. The MI matrices are square adjacency matrices where the MI value corresponds to the weight of interaction for each gene pair. The diagonals are set to zero to ignore self-interactions. The next step is computing row wise ranked versions of these MI matrices in descending order. Here, rank 1 corresponds to the highest mutual information value in a row of the matrix. These ranked matrices will be used in comparing and filtering the networks at the comparison step. Then C3NET is applied to the test and control MI matrices to infer gene networks of direct physical interactions of test and control datasets independently.

We integrated C3NET algorithm with DC3NET package, so we can use all functions of C3NET through DC3NET package’s all in one command. All in one command of DC3NET is as follows:

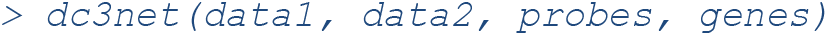

where the first two data set inputs are test and control data sets, e.g. tumor data and healthy data, respectively. Optional input parameters, *method, methodvalue, itnum, rankdif, percentdif, rankdcom*, and *percentcom* are available to control the network inference and decision filtering steps. Furthermore, there are two more parameters, which are *probFiltered* and *visualization*. We recommend enabling *probFiltered* function, since it eliminates the interactions between the probes of the same gene. Visualization function plots the output networks. The visualization parameter takes three values, “0” for disabling the plot, “1” for plotting differential network and “2” for plotting common network. Users can adjust the parameters through command line. The example command above is the simple usage of DC3NET with default parameters.

The parameters that can be set are as follows:

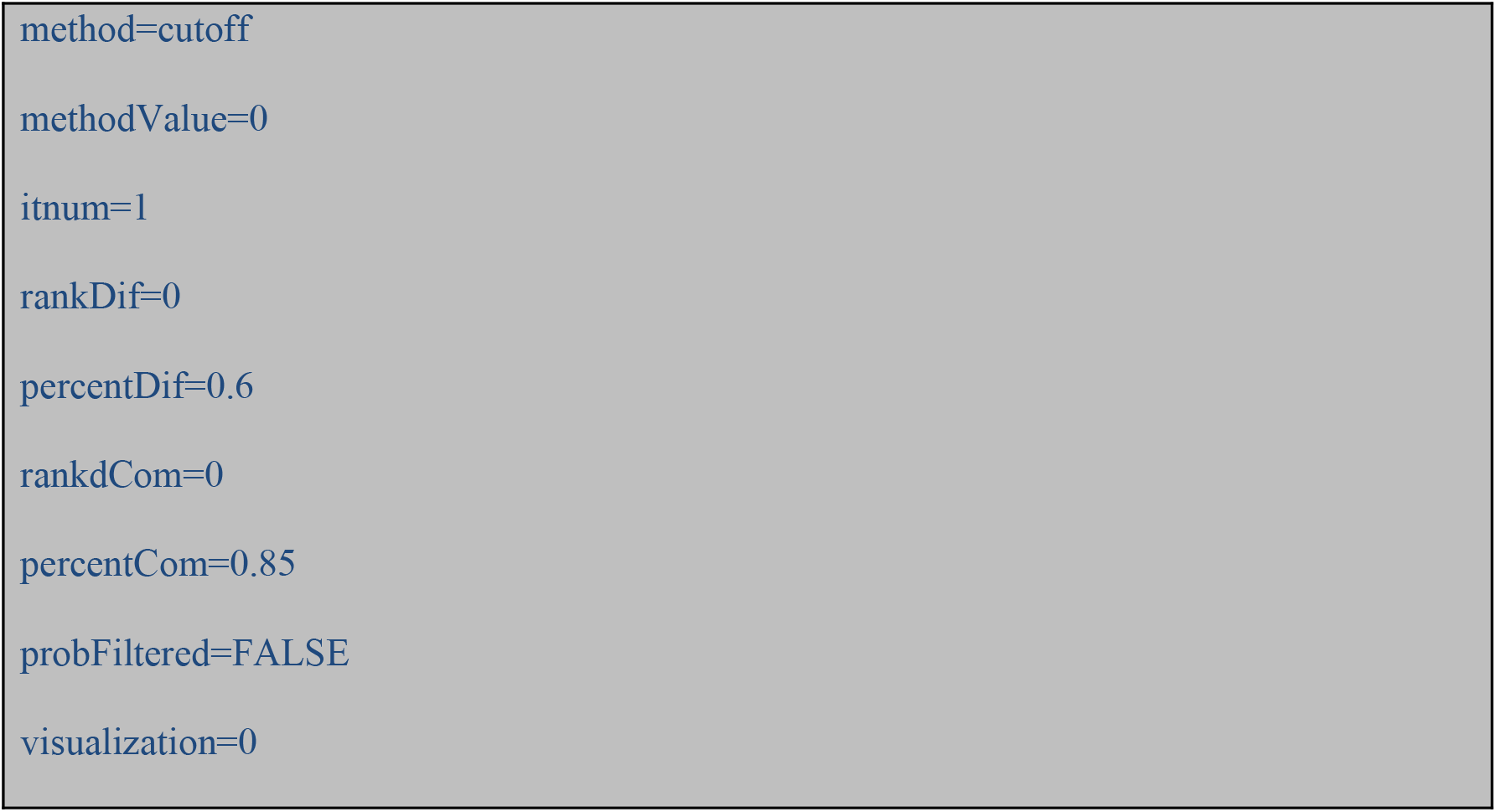

The first three parameters, *method, methodvalue* and *itnum*, are belong to c3net package that are used to eliminate non-significant interactions (Altay and Emmert-Streib, 2010). The available options for *method* parameter are “cutoff”, “justp”, “rank”, “holm”, “hochberg”, “hommel”, “bonferroni”, “BH” and “BY”. *methodValue* and *itnum* parameters are dependent to *method* parameter. If the method is “cutoff”, *methodValue* can be zero or predefined cut-off value. Zero means that mean of upper triangle will be taken as cutoff. If the method is “justp”, *methodValue* and *itnum* (iteration number) parameters are need to be adjusted. For the “justp” method, *methodValue* corresponds to alpha value, e.g. 0.05. If the method is “rank”, methodValue is the rank number. Rank corresponds to the number of interactions that will be taken as significant starting from the highest MI value. The other options, “holm”, “hochberg”, “hommel”, “bonferroni”, “BH” and “BY” are multiple testing correction (MTC) methods. Users can apply MTC methods easily by setting the name of MTC method to the method parameter. If the selected *method* is one of the MTC method, then *methodValue* and *itnum* parameters are need to be adjusted as it were in “justp” method. The default method was set to “cutoff” that uses mean of upper triangle of MI matrices as a significance threshold.

The next four parameters, *rankDif, percentDif, rankdCom* and *percentCom*, are for the comparison step of dc3net. This is the core part of DC3NET that we compare the two networks to find differential networks, *difnet* and *common network*. The order of data1 and data2 is important since the first data set is test network and the second one is control network. If we change the order of input data sets, we find *control difnet*, which consists of interactions that appears only in the control case. It is also crucial since it shows the required interactions of healthy cell.

In the comparison step of DC3NET, there are four conditions that all must be validated at the same time for an edge to be included in *test difnet*. Suppose that we check the potential interaction *geneA* to *geneB* to be included in *difnet* or not. As we stated above, we have been computed row wise ranked versions of the MI matrices in descending order. So we know the rank of interaction *geneA* to *geneB* in *control* MI matrix. The first parameter of DC3NET, *rankdif*, is the predefined cutoff parameter that checks the interaction between *geneA* and *geneB* is one of the top ranked interactions in control MI matrix or not. If the rank of *geneA* and *geneB* in the ranked *control* MI matrix is greater than the predefined cutoff parameter, *rankdif*, then the first condition becomes valid for deciding it as a difnet interaction. *rankdif* parameter can be adjusted to any value between 1 and number of rows of *control* MI matrix. However, if user wants a stricter *difnet*, then *rankdif* parameter needs to be adjusted to a greater value. The second condition is the change in MI value of interaction from *geneA* to *geneB* in the *control* MI matrix. Here, algorithm uses *MIdif* value as the cutoff parameter. *MIdif* is defined as *percentdif* times the maximum MI value of the row of *geneA* in the control MI matrix. Default value for the *percentdif* parameter is 0.6. Depends on strictness of the differential network, user can increase or decrease the second cutoff parameter. The previous two conditions compared the interaction of *geneA* to *geneB* but we also need to compare the interaction of *geneB* to *geneA.* So the algorithm validates the first and second conditions also for the interaction of *geneB* to *geneA.* In this example, if four of the conditions are validated, then DC3NET infer this interaction as in *test difnet* and continue to perform same filtering process for all gene pairs in *test network*.

Lets now start to describe the way that the algorithm infers the *common network. Common network* can be inferred by looking for all the same interactions between test and control network. However, this is a very strict way of inferring common network. So alternatively, one may consider the ranks and MI value decreases in the other data set. More broadly, users may follow the manner of the *difnet* process described above but change the comparison parameter, *rankdif*, from greater to less and for the *percentdif* from less to greater. Additionally, at this time, we only look at one of the two conditions, rather than all the four conditions together, from *geneA* to *geneB* or *geneB* to *geneA.* In the package, *rankdcom*, and *percentcom* parameters correspond to rank difference and MIdif for common network.

Finally, DC3NET infers *difnet* and *common network*, and assign them to the output environment. Furthermore, the package plots the selected inferred network according to the visualization parameter.

### 4. Data structure

#### 4.1. Test data set

Load the example test data set contained in the *dc3net* package:

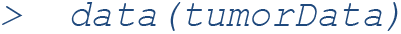

The object tumorData is a tumor data matrix that contains gene expression data for the prostate cancer patients. The sample size is 52. This data was obtained from Broad Institute (Sing *et. al., 2002*). Since the size of the original data set is more than 1 GB, we added randomly selected 500 gene subset of the original prostate cancer data set to the package. This data set can be found in data folder of the package with the filename “tumorData.rda”. The rows of the data set correspond to probes and the columns of the data set correspond to samples.

The dimension of the TEST data set:

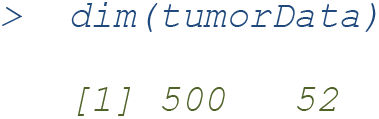

#### 4.2. Control data set

Load the example control data set contained in the *dc3net* package:

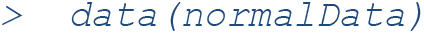

The object normalData is a normal data matrix that contains gene expression data for the prostate cancer patients. The sample size is 50. This data was also obtained from Broad Institute (Sing *et. al., 2002*). The data set can be found in data folder of the package with the filename “normalData.rda”. The rows of the data set correspond to probes and the columns of the data set correspond to samples.

The dimension of the CONTROL data set:

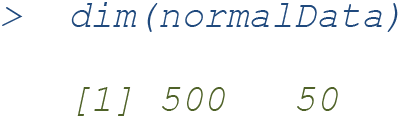

#### 4.3. Probes

Load the gene annotation using one of the input data set. Assign gene annotation to probes parameter:

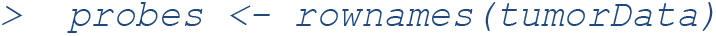

#### 4.4. Gene names

Load the example vector of gene names contained in the *dc3net* package:

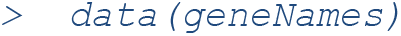

The object geneNames is an R vector that contains the names of the genes inside data sets above. The size of the vector should be equal to the number of rows in your data sets. You should input gene names that correspond to the probes in data sets otherwise the output networks would be produced with probe names. If you can’t obtain the names of the genes, then you should run the *dc3net* command with putting gene annotation to third and fourth parameters e.g. Run dc3net without gene names:

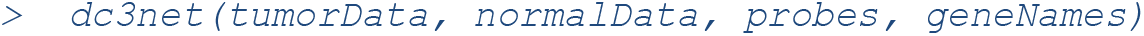

### 5. LnCap example

Here, we show step by step, how to produce the network in Figure 2 of the main paper. In this example, the default parameters were used.

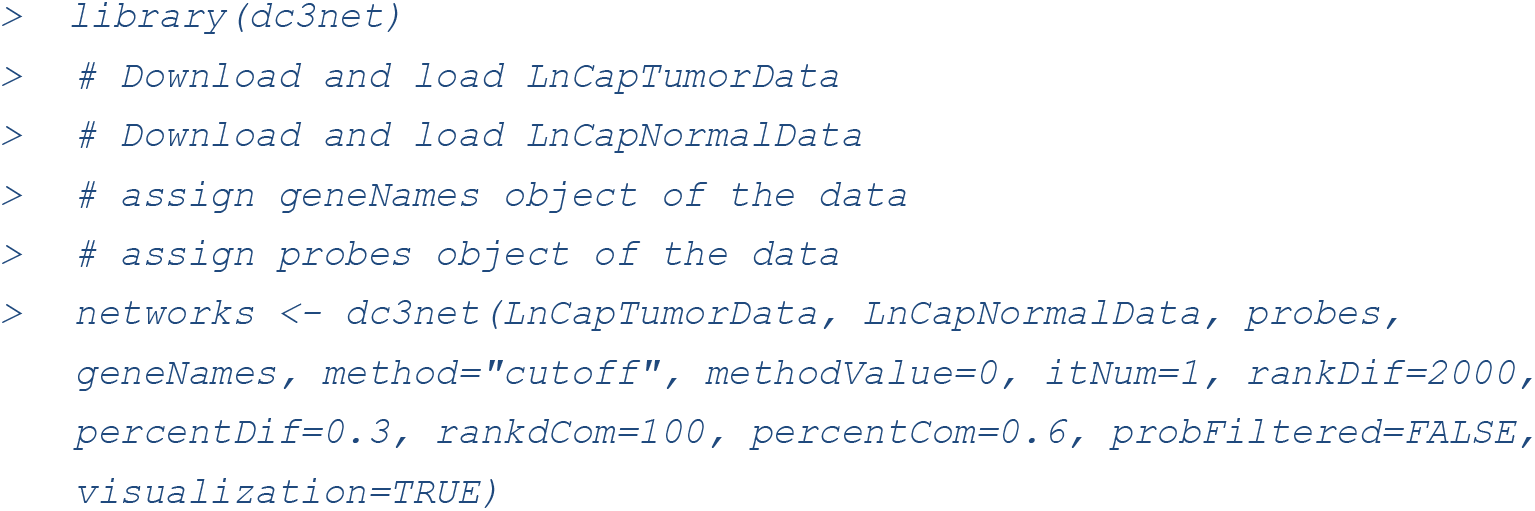

This will compute differential and common networks with the parameters entered through command line. Both commands output differential network and common network tables that can be accessible by:

Differential Network Table:

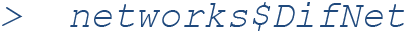

Common Network Table:

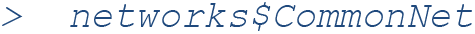

Users can also access to computed mutual information matrices of test and control data by using `networks$mimT` and `networks$mimC`. Thus, users can use this precomputed mutual information matrices on the next run of the algorithm to save time.

The example differential network can be seen in Table 1. In this table, first two columns show the names of the genes that their interactions are in differential or common network. The third column shows the mutual information values between genes. The interactions are sorted according to the mutual information values in descending order. So, the higher rank in the table corresponds to higher interaction. The other columns of the tables can be helpful to advanced users. The fourth column, i, is the row number of the first gene and fifth column is the row number of max partner gene with highest MI value. Sixth and seventh columns are probes of the genes. The next columns are control index, MI rate of the first gene, rank of the first gene, MI rate of the second gene, rank of the second gene in control matrix, respectively.

DC3NET package is designed to inform users when operation continues. Some of the information are the parameters used, the cutoff value computed and dimensions of computed networks. One may easily tune parameters according to this information to obtain better results. The results of DC3NET can be validated through literature using R package, *ganet*. (Altay *et al.*, 2013).

**Table 1.** An example differential network output table of the *dc3net* (The first 10 rows)

| Gene1 | Gene2 | MIval |
| --- | --- | --- |
| LOC729384 | SPRYD5 | 1.67144082 |
| ABCC4 | ACSL3 | 1.53963137 |
| ORM1 | TUBA3E | 1.51528201 |
| SLC41A1 | STK39 | 1.51417854 |
| TLL1 | BCHE | 1.47348983 |
| GLUD1 | SEPP1 | 1.47247699 |
| ACSL3 | IDH1 | 1.45508544 |
| CENPN | ORM1 | 1.45459757 |
| NCAPD3 | CXCR7 | 1.42480939 |
| C11orf92 | UGT2B17 | 1.42160968 |

